# Overactive mitochondrial DNA replisome causes neonatal heart failure via ferroptosis

**DOI:** 10.1101/2022.04.04.485133

**Authors:** Juan C. Landoni, Tuomas Laalo, Steffi Goffart, Riikka Kivelä, Karlo Skube, Anni I. Nieminen, Sara A. Wickström, James Stewart, Anu Suomalainen

## Abstract

Increasing mitochondrial DNA (mtDNA) replication and amount have been proposed as therapeutic approaches for mitochondrial dysfunction, but also as a mechanism of premature aging. We addressed this fascinating paradox by enhancing mtDNA replication via two mechanisms: increasing both mtDNA replication licensing and processivity. We crossed mice overexpressing Twinkle helicase (boosting mtDNA replication initiation) with mtDNA mutator mice (exonuclease-deficient mtDNA replicase, increasing mtDNA mutagenesis and replication processivity). The former model is asymptomatic by two years of age, whereas the latter manifests with progeroid symptoms at six months. Surprisingly, the double transgenics demonstrate postnatally halted growth and devastating cardiomyopathy, fatal within weeks. The mice show high mtDNA replication preventing cardiac maturation and the postnatal shift to oxidative metabolism, causing ferroptotic cardiomyocyte death. Our findings emphasize the critical importance of mtDNA replisome regulation for perinatal cardiac maturation. Furthermore, the data implicate ferroptosis as a cell death mechanism for neonatal mitochondrial cardiomyopathies.

## Introduction

The wide spectrum of clinical manifestations in mitochondrial diseases is unprecedented in the field of medicine. Mitochondrial dysfunction can cause neonatal-onset cardiomyopathies, children’s brain diseases, adult strokes, diabetes, ovarian dysfunction, or parkinsonism ^1^. The mechanisms behind such remarkable diversity of disease signs and symptoms are only starting to be uncovered. Intriguing results from diseases affecting mtDNA maintenance and expression have begun to reveal the mechanisms that could explain such diversity. For example, dysfunction of nuclear-encoded mtDNA replicase (DNA polymerase gamma; POLG) or replicative mtDNA helicase Twinkle cause remodeling of one-carbon (1C) metabolism, affecting the main biosynthetic pathways of the whole cell ^2–5^. 1C-metabolism provides ingredients for growth and repair (nucleotides, phospholipids, amino acids) and serves cells and tissues based on their needs. Furthermore, affected tissues can modify systemic metabolism by secreting metabokines and tuning pathways in distant organs ^3,6–9^. These data reveal the complexity of metabolic signaling in mitochondrial diseases and stress the scarcity of mechanistic knowledge about how different primary defects of mitochondria affect anabolic pathways outside the organelle, in different tissues and ages *in vivo*.

Recent data highlight the cell specificity of mitochondrial manifestations ^10^. The mtDNA “mutator” mice carry an exonuclease-deficient POLG (knock-in of Polg^D257A^) and present mtDNA mutation accumulation alongside a premature aging-like phenotype ^11,12^. These mice have been considered as experimental evidence for the contribution of mtDNA mutagenesis to aging, a hypothesis proposed already 50 years ago ^13,14^. Recently, however, the mutators were shown to also have increased mtDNA replication processivity ^10,15^. Interestingly, in mutator replicating progenitor and stem cells, deoxynucleotide triphosphates were prioritized to mitochondria, starving the nuclear genome of nucleotides and causing a stalled cell cycle at the G1-S phase, slowed-down genomic DNA replication fork progression, and nuclear DNA double-strand breaks ^10^. The progressive manifestation of mutators from six months of age is consistent with stem cell dysfunction: thin skin and hair loss, infertility, and progressive anemia, which becomes fatal at around 12 months ^11,12,16^. The phenotype and nuclear DNA damage mechanism mimics that of other premature aging models ^17^ and suggests a unified mechanism underlying mouse progerias. The only functional defect in postmitotic organ systems in the mutators is late-onset hypertrophic cardiomyopathy ^12^, indicating the sensitivity of the heart to the Polg^D257A^ defect.

Here, we asked whether boosting mtDNA replication frequency in mutators affects the progeroid manifestation. We crossed mutators with mice overexpressing murine Twinkle, showing increased replication licensing and mtDNA copy number ^18,19^. The Twinkle-overexpressors (TwOE) are phenotypically healthy with a normal lifespan of more than two years and develop some respiratory chain deficient muscle fibers at old age ^18,19^. We report here that the combination of a fast mtDNA-replicating mutagenic Polg^D257A^ with TwOE has dramatic synergistic effects in the neonatal heart. The double transgenic mice die shortly after the first week of life because of dilated cardiomyopathy, disrupted cardiac maturation, and ferroptotic death of cardiomyocytes. These results indicate that the regulation of the mtDNA replisome activity is critical for the postnatal metabolic maturation of cardiomyocytes.

## Results

### Increased mtDNA copy number in mtDNA mutators causes growth defects and early death

We crossed the TwOE mice, ubiquitously overexpressing murine Twinkle cDNA (Tyynismaa et al. 2004) with mutator mice carrying a D257A change in POLG (a kind gift from Thomas Prolla; ^11^). The Polg^D257A^ allele was maintained through the paternal line until the last crosses to minimize the accumulation of maternal-inherited mtDNA mutations in the offspring. The crosses resulted in litters with the full array of genotypes: *Polg*^wt/wt^ (WT from now on; carrying some degree of maternal-inherited mtDNA mutations), *Polg*^wt/D257A^ (Polg^Het^), and *Polg*^D257A/D257A^ (Polg^Mut^), and their TwOE experimental counterparts (TwOE, Polg^Het^TwOE, and Polg^Mut^TwOE) (Figure 1A-B). The mice were born phenotypically healthy (Figure 1C), in near-mendelian ratios. The TwOE groups were confirmed to highly express Twinkle mRNA (Figure 1D) leading to a 2-3-fold increase in mtDNA copy number (Figure 1E).

**Figure 1:**
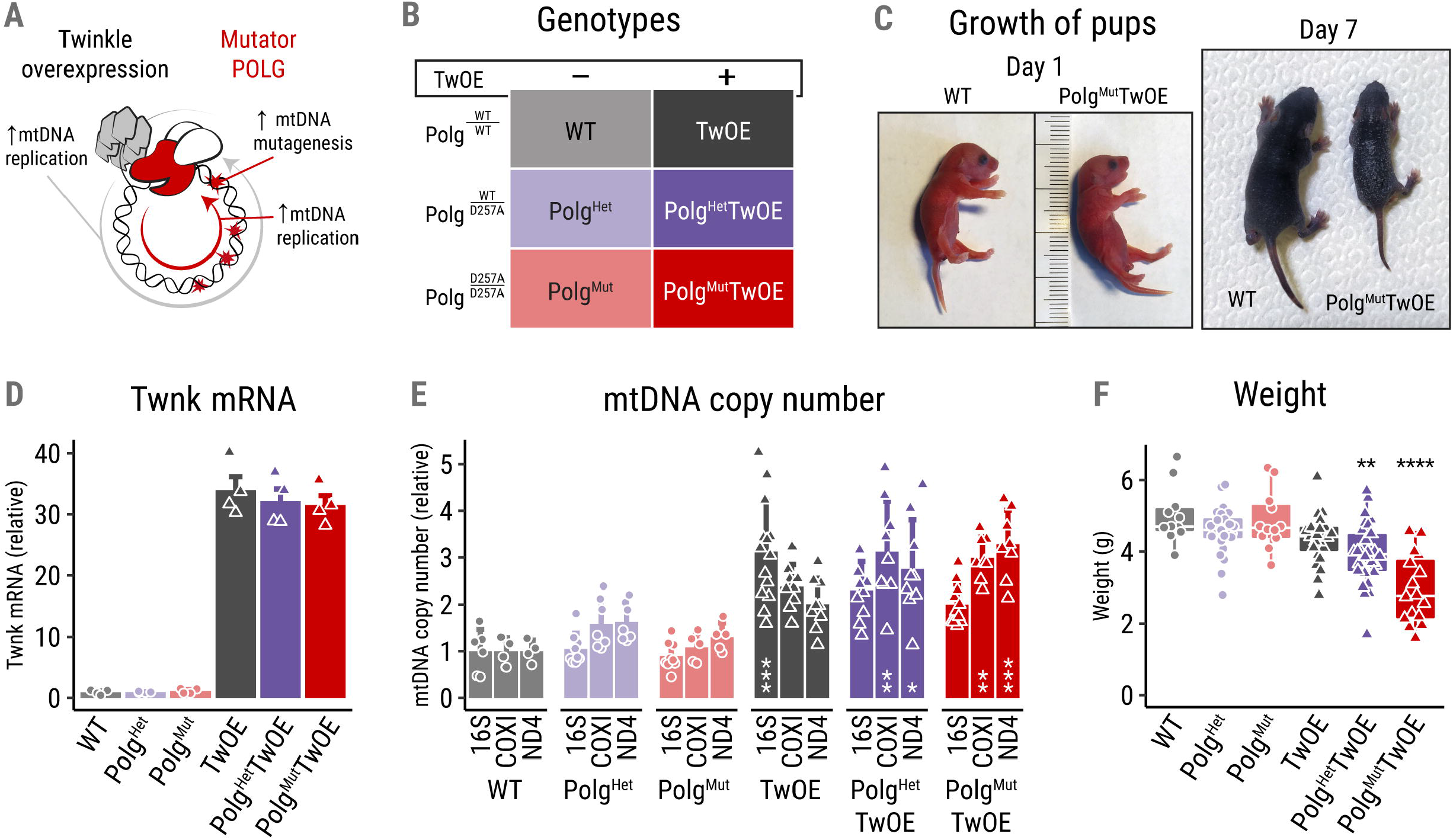
Increased mtDNA copy number in Mutators causes growth defects and early death. (A) Graphical summary of the effects of Twinkle overexpression in mutator *Polg*^wt/D257A^ background for the mitochondrial replication. (B) Array of genotypes in the experimental cohort, with the mutator Polg^D257A^ genotype as rows and TwOE status as columns. (C) Representative images of wild type and Polg^Mut^TwOE mice at postnatal day 1 (P1) and P7. (D) Relative expression of Twnk mRNA in the P1 heart. (E) Relative mtDNA copy number normalized to nuclear gene RBM15, using three different mtDNA sites (genes for 16S rRNA, COXI and ND4) in P7 hearts. (F) Weight of P7 mice in grams. P values from Kruskal-Wallis/Dunn’s test with Bonferroni correction, comparing each group to wildtype controls unless specifically indicated. * (p < 0.05), ** (p < 0.01), *** (p < 0.001), **** (p < 0.0001).

Despite both parental lines with ubiquitous expression of either Polg^D257A^ or TwOE showing late-onset or no visible phenotypes (mutators and TwOE respectively), their offspring developed a striking pathology. The double transgenic mice presented postnatal growth defects already at one week of age (Figure 1C). Lower body weight was associated with TwOE and worsened by the mutant Polg allele amount (Figure 1F). Polg^Mut^TwOE mice started to die at postnatal day 9, with less than half reaching three weeks of age. These results suggest that licensing more mtDNA molecules to enter replication, combined with a fast and mutagenic POLG is highly deleterious in postnatal life.

### Enhanced mtDNA replication in mice carrying at least one *Polg*^D257A^ allele associates with neonatal heart failure and multi-organ damage

The surprising neonatal synthetic lethality of TwOE and Polg^D257A^ prompted us to search for the cause of death. We found Polg^Het^TwOE and Polg^Mut^TwOE mice to develop a rapidly progressive cardiac enlargement by one week of age, despite presenting no apparent disease signs at birth (Figure 2A). The histological inspection showed interstitial fibrosis, vacuolation, and cardiomyocyte degeneration and lysis, leading to large infarcted regions (Figure 2B-D), with ∼60% penetrance of the heart phenotype in Polg^Het^TwOE and full penetrance in Polg^Mut^TwOE. The hearts of the Polg^Mut^TwOE mice showed enlarged cardiomyocytes with disrupted mitochondrial ultrastructure and large and less abundant nuclei than in WT hearts (Figure 2D-F), the latter suggesting defective nuclear polyploidization that typically occurs during the first week of life ^20^. Echocardiography at postnatal day 7 (P7) confirmed the severe heart failure in the double transgenic groups, with decreased ejection fraction, large left ventricle volume, and thin ventricle walls, characteristics of dilated cardiomyopathy (Figure 2G-J).

**Figure 2:**
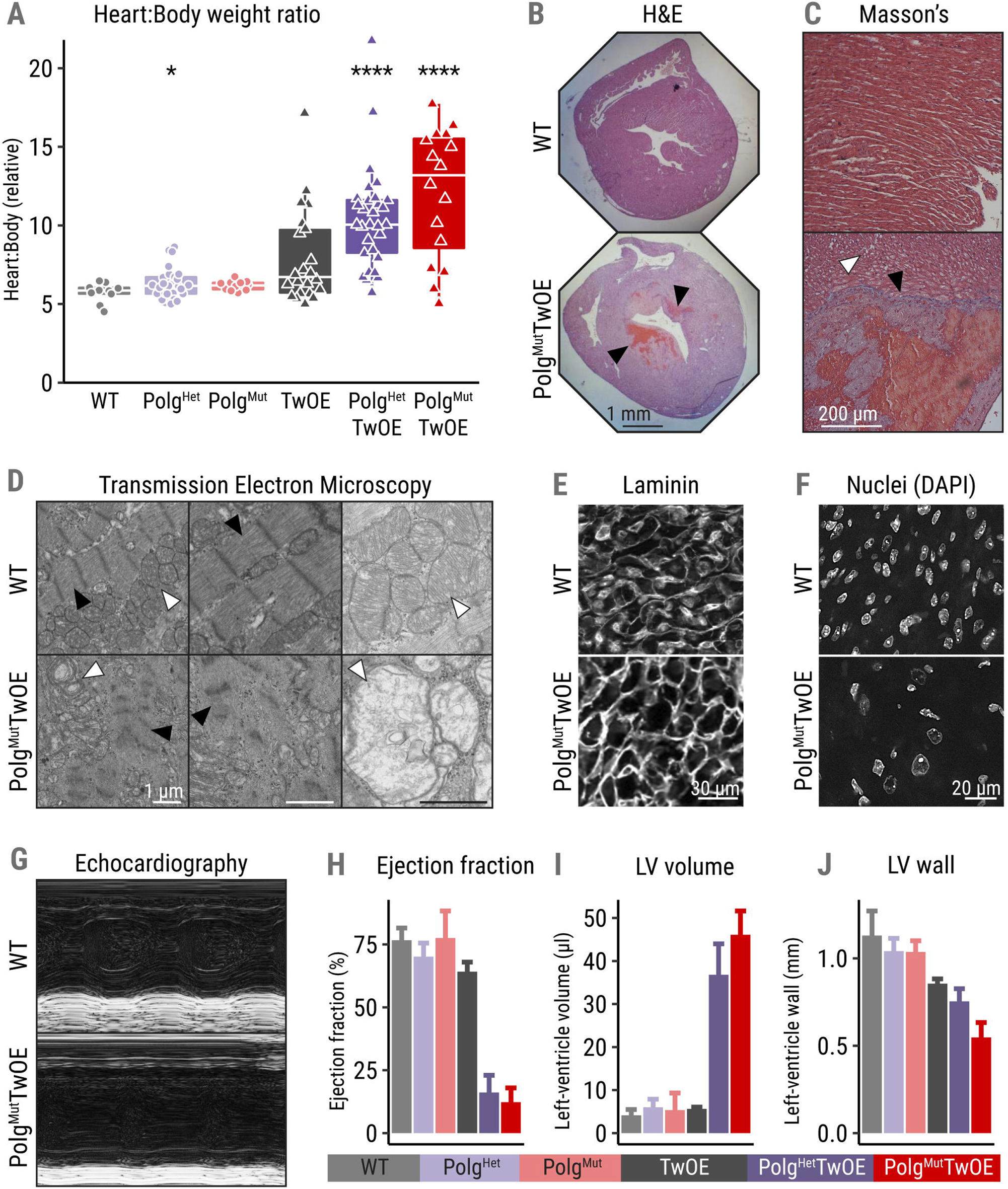
Enhanced mtDNA replication in mice carrying one or two mutator Polg alleles associates with neonatal heart failure and multi-organ damage. (A) Heart to body weight ratio at P7. P values from Kruskal-Wallis/Dunn’s test with Bonferroni correction, comparing each group to wildtype controls. * (p < 0.05), ** (p < 0.01), *** (p < 0.001), **** (p < 0.0001). (B-D) Representative images of P7 heart characterization: (C) Infarction-like pathology (black arrowheads); hematoxylin/eosin staining. (C) Myocardial disarray, vacuolation (white arrowheads) and interstitial fibrosis (black arrowheads); Masson’s trichrome staining. (D) Lysis of the myofibrillar structure (black arrowheads), and the disruption of mitochondrial membrane ultrastructure (white arrowheads); transmission electron micrograph. (E) Increase in cardiomyocyte size; P7 hearts, representative image of laminin immunostaining; (F) Enlarged and less abundant nuclei; representative image; DAPI staining. (G-J) Dilated cardiomyopathy and severe heart failure in Polg^Het^TwOE and Polg^Mut^TwOE; echocardiography, quantified indicators.

At P7, the liver was fibrotic with severe fat accumulation, pointing to decreased blood flow via the inferior *vena cava* to the right atrium of the heart (Figure S1A). The fat accumulation could also be a result of the inability to oxidize fat. The lungs had disrupted alveolar structure (Figure S1B), suggesting pulmonary hypertension due to failing pulmonary artery bloodstream to the heart. Neither organ presented significant changes in mtDNA copy number at P7 or signs of damage at P1 (Figure S1C), pointing to the heart as the primary failing organ. We also assessed the tissues typically affected in the progeroid adult mutator mice (hemoglobin, hair growth). While blood hemoglobin and glucose concentrations showed a mild decrease (Figure S1D, E), the replicating cells such as testes, skin epidermis and hair follicles appeared unaffected, without major changes in proliferation or DNA damage at the early age of P7 (Figure S1F).

In summary, our results indicate that increased mtDNA replication in mice with mtDNA mutagenesis is not tolerated in a postnatal heart. Together, TwOE and Polg^D257A^ trigger heart failure and are synthetic lethal for perinatal heart, with devastating multiorgan consequences.

### Enhanced mtDNA replication exacerbates the Polg^D257A^ effects for mtDNA stability

POLG and Twinkle co-operate in the mtDNA replication fork, and therefore we asked whether mtDNA rearrangements or mutagenesis explained the dramatic cardiac manifestation. The size of mtDNA nucleoids/nucleoid clusters was increased in both cardiac tissue and cultured mouse embryonic fibroblasts (MEFs) of TwOE mouse groups (Figure 3A-B; S1G). In TwOE and Polg^Mut^TwOE MEFs, mtDNA replication frequency was increased, as measured by EdU incorporation (Figure 3C). Surprisingly, the nucleoid size difference was exclusive to actively replicating nucleoids (Figure 3B), suggesting that more mtDNA molecules associated with one nucleoid underwent replication than in wild types, and the newly synthesized molecules were segregated after completion. We then performed deep-sequencing of mtDNA to study whether increased replication had resulted in the accumulation of specific mutations that would explain the severe phenotype. Polg^Het^TwOE and Polg^Mut^TwOE presented increased mutation loads when compared with their non-TwOE controls, consistent with more frequent mutagenic events due to augmented mtDNA replication (Figure 3D). However, the detected mutations remained at very low heteroplasmy levels (<10%) and were similar across the genotypes, not explaining the phenotype (Figure 3E). When separated by type of misincorporation, the vast majority of mutations were T or A misincorporations (C:G>T:A and A:T>T:A), consistent with the known behavior of Polg^D257A 21^ (Figure 3F).

**Figure 3:**
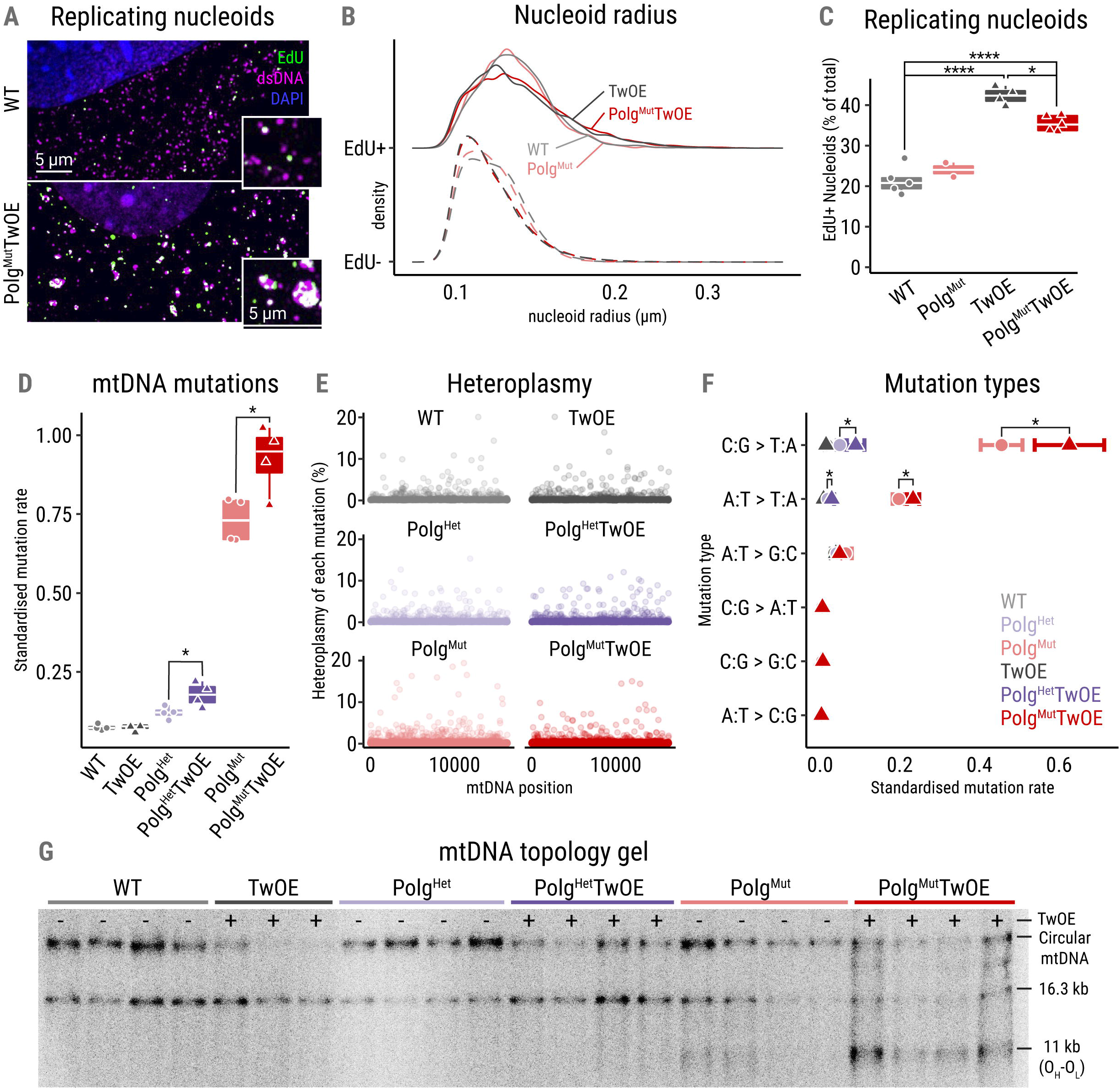
Enhanced mtDNA replication exacerbates the consequences of Mutator Polg in mtDNA stability. (A) Replicating nucleoids; representative fluorescent microscope image of embryonic fibroblasts; double-stranded DNA antibody (dsDNA, nucleus and nucleoids, magenta), EdU (replicating mtDNA, green) and DAPI (nucleus, blue). (B) Nucleoid radius distribution by genotype and by replication status (EdU+ as full lines, EdU- as dashed lines). (C) Quantification of nucleoids in replication (EdU+) compared to total nucleoid number. P values from ANOVA/Tukey’s HSD test; * (p < 0.05), ** (p < 0.01), *** (p < 0.001), **** (p < 0.0001). (D) Mutation rate in P7 heart mtDNA; next-generation sequencing, total number of mtDNA mutations detected standardized to the occurrence of each base. P values from a two-tailed t-test contrasting groups with identical POLG genotype. * (p < 0.05), ** (p < 0.01), *** (p < 0.001). (E) Heteroplasmy of each point mutation detected across the mtDNA sequence, as a percentage of the total reads for that locus. (F) Mean standardized mutation rate by substitution type, with SEM. P values as in (D). (G) MtDNA topology; Southern hybridization analysis showing the different topological species of mtDNA present in the sample.

MtDNA topology analysis revealed a remarkable increase of abnormal mtDNA forms, with a decrease in the relaxed circular forms in mice overexpressing Twinkle, and a high amount of linear mtDNA fragments spanning between the two replication origins of mtDNA were especially enriched in the Polg^Mut^TwOE (Figure 3G; S1H-I). However, again, the presence of the abnormal mtDNA topologies did not correlate with disease manifestation.

Together, these results indicate that enhancing mtDNA entry to replication exacerbates the consequences for mtDNA integrity previously reported to occur in mutator mice. However, these considerable changes still do not explain the severity of the heart failure, as a single Polg^D257A^ allele (in Polg^Het^TwOE) showed no measurable consequences for mtDNA integrity, while both Polg^Het^TwOE and Polg^Mut^TwOE presented with heart failure.

### Disrupted shift to oxidative metabolism in double transgenics: perinatal integrated mitochondrial stress response and metabolic remodeling with coenzyme-A deficiency

At P7, the hearts of the double transgenic mice were already severely affected. Therefore, to explore the pathophysiology leading to cardiac dysfunction events after birth, we performed proteomic and metabolomic analyses from P1 mouse hearts.

The proteomics analysis detected 2423 proteins significantly changed (Qval < 0.01) between Polg^Mut^TwOE and WT after multiple testing correction (Figure 4A-B & S2A). The analysis revealed the typical signature of the mitochondrial integrated stress response (ISRmt), including increased expression of *de novo* serine biosynthesis enzymes (PHGDH, PSPH, PSAT1), asparagine synthetase (ASNS), the mitochondrial folate cycle (MTHFD2, MTHFD1L), and proline metabolism (PYCR1). In addition, cytosolic aminoacyl tRNA synthetases and amino acid transporters were upregulated as well as cytoplasmic ribosomal subunits and their assembly factors, all suggesting induction of cytoplasmic translation and amino acid metabolism, while mitochondrial translation machinery components were low in amounts. Enrichment for the mitochondrial proteome using the MitoCarta3.0 database ^22^ indicated that almost 50% of all mitochondrial proteins were significantly changed in Polg^Mut^TwOE mice, the majority being downregulated (Figure 4C). Especially, the nuclear-encoded subunits of the respiratory chain enzymes were reduced, most significantly those encoding complexes I, III and IV (Figure 4D), indicating downregulation of the oxidative phosphorylation machinery. Additionally, the biosynthesis of coenzyme Q, a key component of the respiratory chain and the cellular antioxidant metabolism, was downregulated. Also, genes involved in heart contraction and cytoskeleton/myofibrillar composition were downregulated, while profibrotic secreted factor CCN2 was increased. As a whole, markers of cardiomyopathy were induced, as classified by the KEGG database ^23^, suggesting that the cardiac damage signaling is either acutely induced within one day of postnatal life, or taking place already neonatally, before the onset of visible damage. We did not detect changes in the nucleotide metabolic pathway proteins or nuclear DNA repair proteins, as in adult stem cells, indicating the specificity of the changes to the neonatal heart.

**Figure 4:**
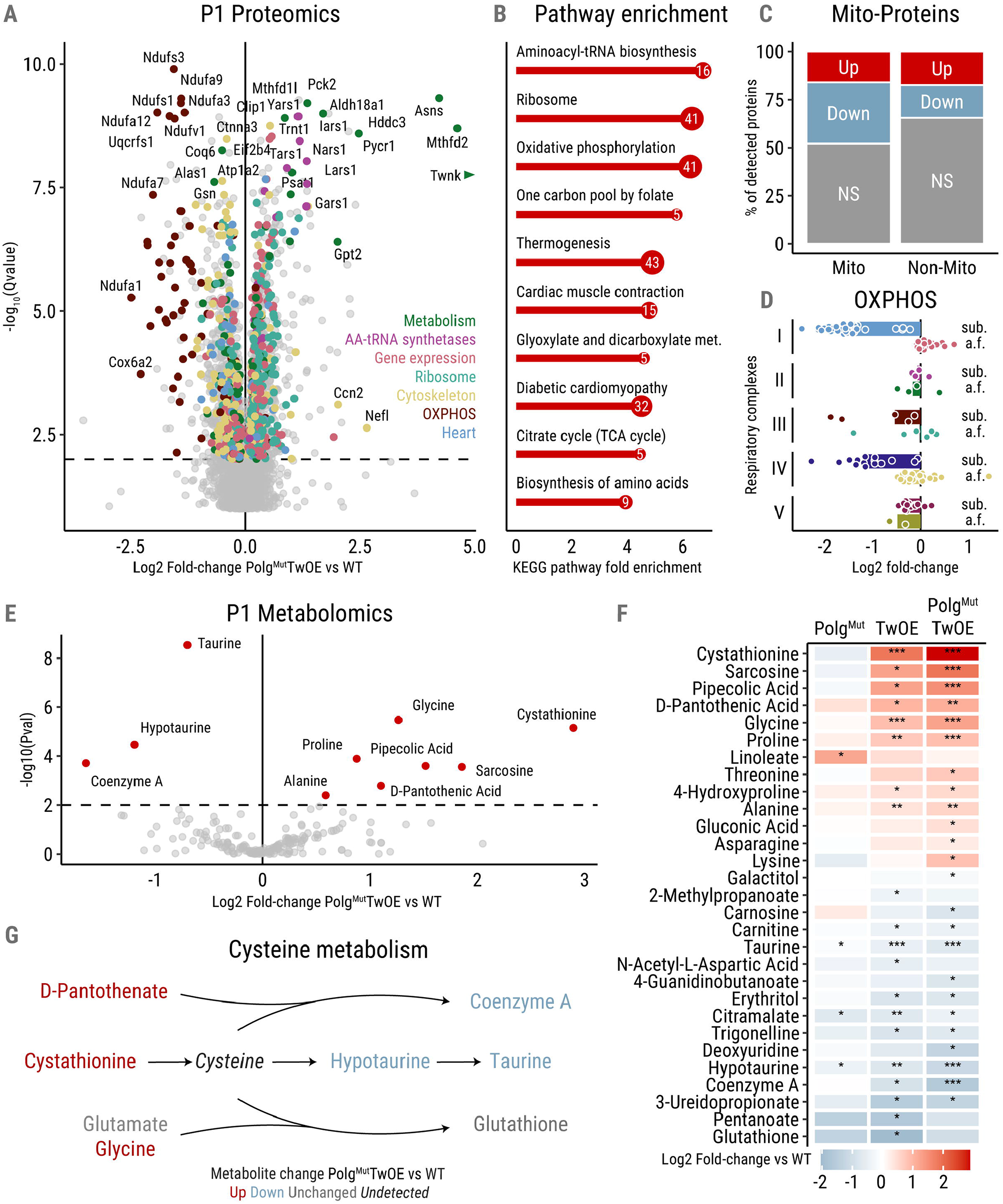
Multi-omic approaches reveal perinatal ISRmt induction, cysteine metabolic defects and large-scale mitochondrial and metabolic rewiring. (A) Volcano plot depicting the fold-change and statistical significance of the proteomic changes between WT and Polg^Mut^TwOE groups. The dotted line indicates the significance threshold at Q < 0.01 and changed hits are categorized into groups. Twnk arrowhead indicates the position of the cropped datapoint. (B) Enriched KEGG pathways among the proteomic hits, fold enrichment on the x-axis and log10 p-value in the lollipop plot. (C) Percentage of changed hits classified as mitochondrial and not according to MitoCarta3.0. (D) Changes in respiratory chain subunits (sub.) and assembly factors (a.f.) by respiratory complex. (E) Volcano plot depicting the fold-change and statistical significance of the metabolite’s change between WT and Polg^Mut^TwOE groups. The dotted line indicates the significance threshold at p < 0.01 and changed hits are highlighted in red and named. (F) Heatmap displaying the Log2 fold change of the significantly affected metabolites across all measured genotypes. Raw P-values from a t-test to the WT control as stars: * (p < 0.05), ** (p < 0.01), *** (p < 0.001). (G) Simplified cysteine metabolic pathway integrating and depicting some of the significant changes from the metabolomic analysis.

A targeted metabolomic analysis of 196 metabolites of P1 hearts showed findings that converged especially in cysteine metabolism. Remarkably, coenzyme A (CoA), the key regulator of lipid synthesis and oxidation, was found to be decreased both in TwOE and Polg^Mut^TwOE mice (Figure 4E-F). Although our metabolomics setup was not able to detect cysteine directly, the accumulation of cystathionine (cysteine precursor) and pantothenate (CoA precursor), together with decreased hypotaurine, taurine and glutathione (significant in TwOE) all suggest a deficiency of cysteine derivatives (pathway overview in Figure 4G).

To clarify whether the changes were acute or already initiated during fetal life, we investigated the transcriptomic landscape in E16.5 mice. The analysis revealed 174 significantly changed transcripts between Polg^Mut^TwOE and WT (Figure 5A), a much more modest figure than the proteomics of P1 mice, especially when considering the 3-fold larger set of detected genes. Interestingly, the main pathways induced were ISRmt and the cytosolic tRNA synthetases, with ATF5 as the strongly upregulated effector of the response (Figure 5A-B). Most of the downregulated genes participated in the extracellular collagen matrix maintenance and heart contraction, indicating that already during embryogenesis, the hearts were undergoing changes in myocardial structure.

**Figure 5:**
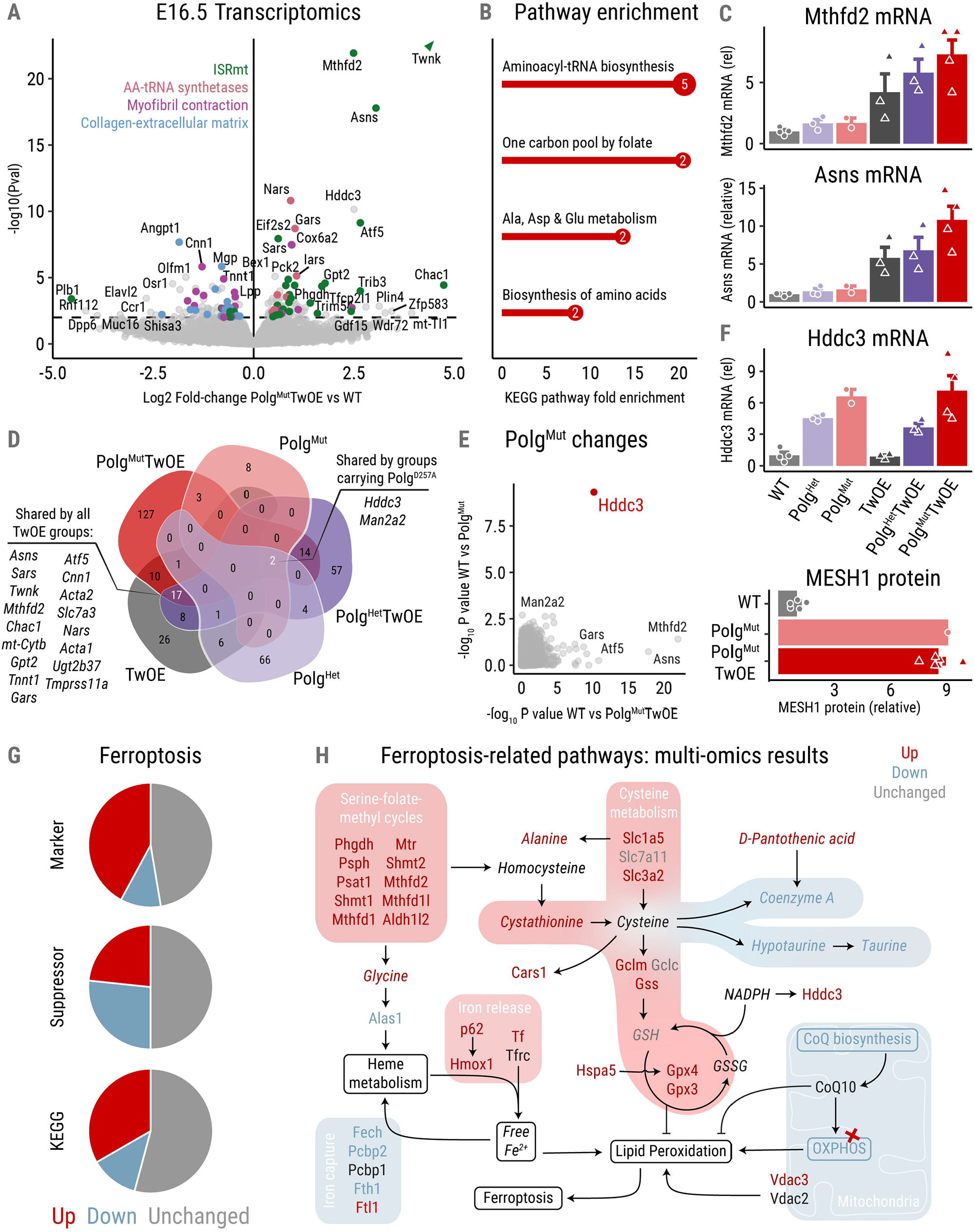
Perinatal induction of the mitochondrial stress response and pro-ferroptotic signaling fatally disrupts cardiomyocyte metabolic maturation. (A) Volcano plot depicting the fold-change and statistical significance of the transcriptomic changes between WT and Polg^Mut^TwOE groups. The dotted line marks the significance threshold at p < 0.01 and changed hits are categorized into groups. Twnk arrowhead indicates the position of the cropped datapoint. (B) Enriched KEGG pathways among the transcriptomic hits, fold enrichment on the x-axis and log10 p-value in the lollipop plot. (C) Example plots of the landscape of mRNA change across all genotypes in ISRmt genes. (D) Venn diagram of the overlap in significantly changed genes across the genotypes. Interesting groups are highlighted and detailed as lists. (E) Scatterplot comparing the statistical significance of the change in Mut and Polg^Mut^TwOE groups (against WT), resulting in *Hddc3* as the sole shared highly significant gene. (F) Relative amounts of Hddc3/MESH1 mRNA and protein in different genotypes, showing dose-responsiveness to Polg^D257A^ alleles. (G) Pie charts showing the proportion of proteins marked as ferroptosis marker or suppressor (FerrDb), or in the KEGG ferroptosis pathway, found significantly (Qval < 0.01) upregulated (red) or downregulated (blue) or non-significantly changed (grey) in Polg^Mut^TwOE. (H) Main pathways involved in ferroptosis, overlaid with the change detected by proteomics (normal font) and metabolomics (italics) with the same color scheme as (G) and black as not measured/detected.

**Figure 6:**
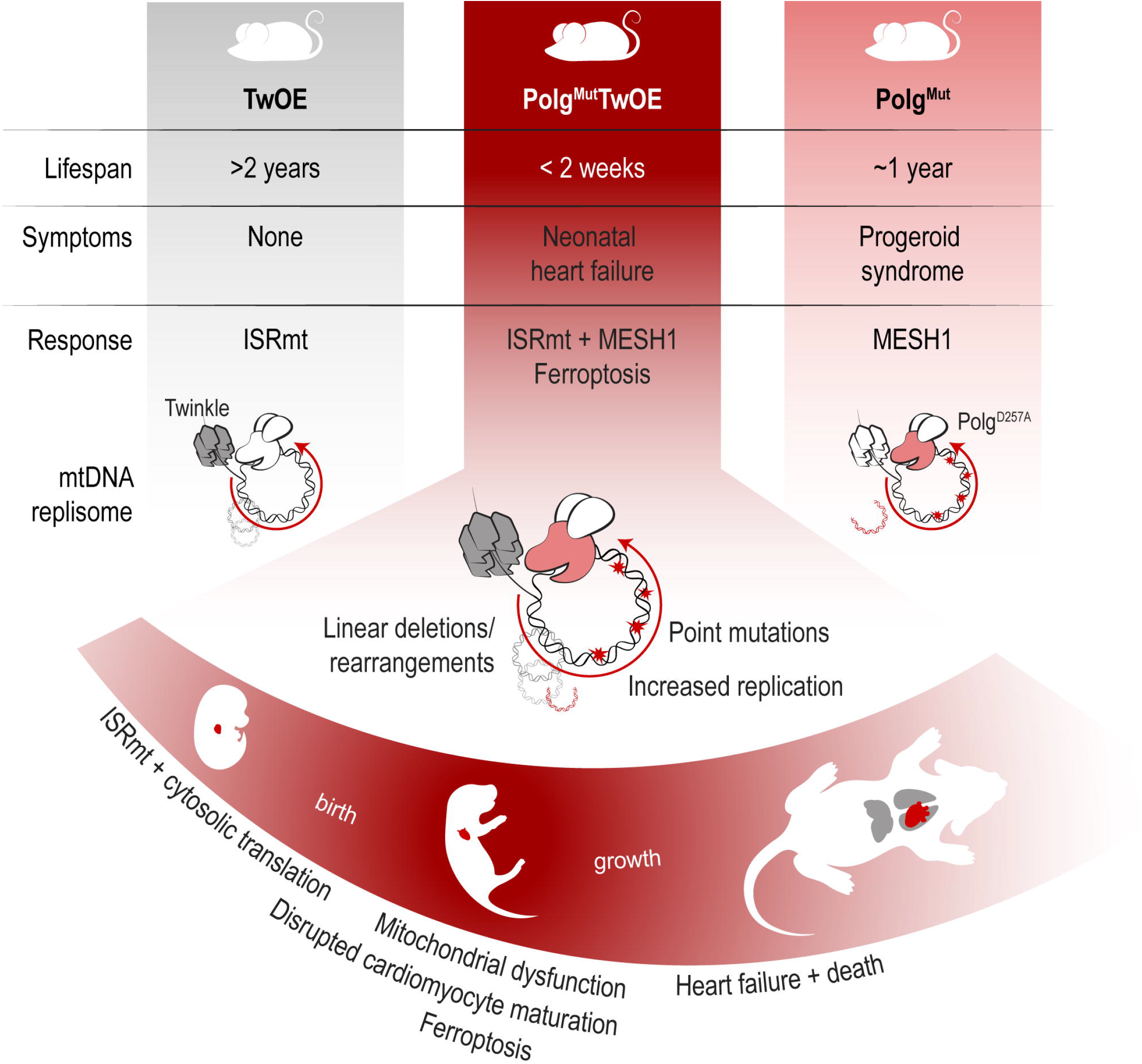
Overactive mitochondrial DNA replisome causes neonatal heart failure via ferroptosis. Summary of the findings, contrasting the symptomatology and pathomechanism of the parental/control mouse lines with the severe early-onset pathology of their combination. The graphical depiction of the mtDNA replisome highlights the affected proteins in each model, as well as their key effects on mtDNA replication and integrity. Finally, the perinatal timeline indicates the main affected organs in each stage of maturation and the critical processes leading up to the early death of the double transgenic mice.

### Ferroptotic cell death induced by mitochondrial dysfunction in the heart

Many transcriptomic changes correlated strongly to the severity and incidence of the cardiac failure, exclusive to mice that overexpressed Twinkle in mutator background, dose-responsibly exacerbated by the Polg^D257A^ alleles (Figure 5C-D). Even TwOE alone, if born from a mutator heterozygote female, induced a mild ISRmt in the heart (Fig 5D), potentially to increase nucleotide synthesis as a response to increased mtDNA replication. Therefore, we next sought to identify factors that would uniquely and dose-responsively respond to Polg^D257A^ genotype but not to TwOE as potential candidates to pathologically synergize in the heart. Comparing the significantly changed transcripts in both Polg^Mut^ and Polg^Mut^TwOE mice at E16.5, the top-most hit was *Hddc3* (Figure 5E), which proportionally correlated with Polg^D257A^ allele number (Figure 5F). *Hddc3* encodes for the NADPH phosphatase MESH1 which regulates ferroptosis ^24^, a recently described process of non-apoptotic cell death ^25^.

*Hddc3*/Mesh1 was also strongly upregulated at the protein level (Figure 5F). We then filtered the significantly changed proteins of P1 mice using the manually curated ferroptosis database FerrDb ^26^ and KEGG pathway ^27^. Indeed, about half of the FerrDb markers and KEGG ferroptosis pathway proteins were changed (20 out of 38 and 11 out of 24 respectively), typically upregulated, and half of the ferroptosis suppressors were significantly changed (15/30), most of which downregulated (Figure 5G). Some of the key changes included the induction of the central ferroptosis regulator GPX4, dysregulation of iron metabolism in the direction of iron release (notably the induction of heme catabolism by HMOX1 and inhibition of heme synthesis and iron uptake by ALAS1 and FECH1), as well as disruptions of the mitochondrial respiratory chain and coenzyme Q biosynthesis. These data fill the criteria of ferroptotic pathway induction and suggest that the mutator mice are primed to ferroptosis, which then in our current double transgenics was triggered by TwOE. The summary of the remarkable induction of the canonical ferroptotic processes revealed by our multi-omics approach is presented in Figure 5H.

## Discussion

We show here that increased mtDNA replication is deleterious for perinatal heart development if co-occurring with mtDNA mutations. The synergistic pathogenic effect of the two enzymes at the mtDNA replication fork –Polg^D257A^ and Twinkle- was surprising, because the parental mouse lines show late-onset or no symptoms [mutators, after 6 months of age ^11,12^; TwOE mice, healthy >2 years of age ^18,19^]. As the overexpressed Twinkle was wildtype, the only mutant in the double transgenics is the exonuclease-deficient POLG. This deficiency results in the accumulation of random mtDNA mutations, but also “removes the brakes” from the accurate DNA polymerase, leading to increased replication frequency ^10^. A further boost of the replication frequency, by the Polg^Mut^TwOE combination, allows for normal fetal development but is synthetic lethal for the neonatal heart, causing the hepato-cardio-pulmonary system to collapse within a week. Our multi-omic analysis indicates a major remodeling of one-carbon metabolism with typical components of ISRmt already at E16.5. These changes do not challenge normal development. Neonatally, however, the double transgenic mice are unable to undergo the typical steps of neonatal cardiac maturation: to shift to oxidative metabolism and use of lipids as a fuel, synthesize CoA and undergo nuclear polyploidization. The hearts show a deficiency of cysteine derivatives, while paradoxically boosting components of cytoplasmic translation. Our evidence highlights the remarkable importance of mtDNA replisome control for tissue homeostasis. Losing that control leads to neonatal dilated cardiomyopathy with lytic death of cardiomyocytes and a strong signature of ferroptosis.

Counterintuitively to our current results, mtDNA copy number regulation has been explored as a potential way to treat ischemic heart failure ^28–31^ and mitochondrial diseases with mtDNA mutations ^32–34^. Overexpression of TFAM, the mtDNA packaging protein and transcription factor, rescued mutator mouse testicular degeneration and prevented nuclear DNA damage in mutator stem cells ^10,34^, suggesting that increased packaging slows down mtDNA replication by Polg^D257A 10,19,35,36^. Furthermore, TwOE using the mouse line of the current study was shown to be protective against genetic and ischemic events of the heart ^28–31^. Indeed, TwOE crossed to SOD2+/- mice prevented the progression of hypertrophic cardiomyopathy of the latter. In these mice, despite the protective effects, TwOE did not prevent the generation of large-scale mtDNA rearrangements, agreeing with our data that mtDNA topology changes were unrelated to the pathology. A combined expression of Twinkle and TFAM resulted in enlargement of nucleoids, multiple mtDNA deletions, reduced transcription and late-onset mild respiratory chain dysfunction ^19^, but the mice remained healthy and lived a normal lifespan. These data indicate that increased licensing to mtDNA replication by TwOE can be curative. However, when combined with accelerated processivity in the background of mtDNA mutations, the effects are deleterious in early life. Our evidence highlights the need for caution, especially for approaches aiming to boost mtDNA replication in diseases with mtDNA mutations.

The first week of postnatal life is crucial for heart development. The heart undergoes a rapid shift from fetal glycolytic glucose-dependent metabolism to ketone bodies and lipid oxidation with increased tissue oxygen levels ^37^ also repressing the mitochondrial compartment of 1C metabolism ^38^. Oxidation of the new fuels requires an active mitochondrial respiratory chain, the function of which promotes terminal differentiation. During the first week, the heart is still able to regenerate, and cardiomyocytes undergo nuclear division, resulting in mature polyploid cells ^20,39^. Of biosynthetic growth-promoting metabolism, a previous report identified cysteine, taurine, methionine and glutathione metabolism as one of the most enriched pathways during the first postnatal week, alongside an increase in CoA and its biosynthesis intermediate phosphopantetheine ^37^. Remarkably, in our Polg^D257A^TwOE mice, OXPHOS remains uninduced, the fetal pattern of 1C-metabolism keeps being high, and nuclear morphology is abnormal and less abundant than in WTs. The results indicate that overactive mtDNA replisome results in a complete inability to shift to postnatal cardiac maturation.

The metabolic remodeling in the Polg^Mut^TwOE mice offered several significantly changed pathways that can explain the early lethality converging in ferroptosis. 1) Imbalance of 1C-flux to transsulfuration pathway that synthesizes cysteine, glutathione and taurine, all essential for cardiac homeostasis ^37,40,41^; 2) Mitochondrial respiratory chain deficiency and consequent redox imbalance, together with the downregulation of coenzyme Q biosynthesis, a component of respiratory chain and a factor inhibiting lipid peroxidation ^42,43^; 3) Depletion of CoA, essential for lipid synthesis and oxidation; 4) Induction of amino acid synthesis and cytosolic translation, the main user of ATP in cells, in the context of unsuccessful OXPHOS activation; 5) Dysregulation of iron and heme metabolism, including the induction of the iron-releasing and ferroptotic enzyme HMOX1, and downregulating heme synthesis and ferrous reuptake enzymes ALAS1 and FECH1; 6) Induction of canonical ferroptotic responses mediated by GPX4, CHAC1, VDAC3, HMOX1 and NRF2 target genes ^44^, as well as *Hddc3*/MESH1 ^24^. The lack of cardiac nucleotide synthesis or DNA-repair enzyme changes, or genomic DNA breaks, as previously reported in stem cells ^10^, highlights the tissue- and age-specificity of the sensitivity points in metabolism. The data tightly connect mtDNA replisome signaling to cellular amino acid, redox, lipid, and iron homeostasis.

Ferroptosis is a recently discovered form of regulated cell death, characterized by the iron-dependent accumulation of lipid reactive oxygen species exceeding the cellular antioxidant capacity and causing the fatal collapse of redox homeostasis ^25,45,46^. While OXPHOS function is required for the induction of cysteine deficiency-mediated ferroptosis ^47^, the role of primary mitochondrial dysfunction in triggering ferroptosis has been under debate ^48,49^. Recent reports linked drug-induced mtDNA damage to autophagy-dependent ferroptosis in pancreatic cancer ^50^. Also, mtDNA depletion in hepatic organoids led to ferroptosis ^51^. Furthermore, cardiac ischemia/reperfusion and chemotherapy-induced cardiomyopathy have been associated with mitochondrial membrane peroxidation-mediated ferroptosis ^52^. The upregulation of Hmox1 and a mitochondria-dependent cascade were the main effectors of the cardiac damage ^52,53^. Our data joins the body of evidence associating mitochondrial dysfunction with ferroptosis and cardiac failure, highlighting that mtDNA replisome integrity may be a critical signal for ferroptosis and suggesting a novel role for ferroptosis in early heart maturation and disease.

Mitochondrial cardiomyopathies are devastating forms of mitochondrial disease, typically lethal at an early age and most often caused by defects in mtDNA expression, especially translation ^54^. Our data demonstrate that tight regulation of mtDNA amount and replication is critical for perinatal heart development, and highlights the importance of mtDNA homeostasis in healthy cardiomyocyte maturation, along with a potential role for primary mtDNA maintenance in ferroptotic death signaling.

## Acknowledgments

The authors wish to thank Markus Innilä, Tuula Manninen, Sonja Jansson, Babette Hollmann, Satu Malinen, and Maria Arraño de Kivikko for their technical contributions and expertise; Amy Platt, Aurora Hämälainen and the team of animal caretakers are thanked for their excellent handling and flexibility. We want to thank Howard Jacobs for the enriching scientific discussion, and Olesia Ignatenko, Gulayse Ince Dunn, Rocio Maldonado, Swagat Pradhan and Saara Forsström for their support and technical contributions. We thank the core facilities that enabled the performance of this work: the Electron Microscopy Unit at the Institute of Biotechnology, the Biomedicum Functional Genomics Unit, the FIMM Metabolomics Unit (funded by Biocenter Finland and HiLIFE), the Turku Proteomics Facility at Biocenter Finland, and Franziska Metge of the Bioinformatics Core facility at MPI-AGE, Cologne, Germany. We further wish to acknowledge the Sigrid Jusélius Foundation, Academy of Finland, Helsinki University Hospital (for A.S.), Maud Kuistila Foundation, Päivikki ja Sakari Sohlberg Foundation, Biomedicum Helsinki Foundation, and the University of Helsinki Funds (J.C.L.) for funding the study.

## Author Contributions

Conceptualization: JCL, AS; Investigation: JCL, TL, SG, RK, KS, JS, AN, SW; Funding acquisition: AS, JCL; Supervision: AS; Writing – original draft: JCL, AS; Writing – review & editing: all authors.

## Materials and Methods

### Mouse models and cell culture

The mouse strains used were the Mutator mice, with a knock-in D257A mutation in DNA polymerase gamma ^11^, and the Twinkle overexpressor mice, carrying a Twnk cDNA transgene ^18^. The experimental group consists of the different genotype combinations arising from the cross between those two strains (Figure 1B), obtained by first crossing *Polg*^wt/D257A^ males with TwOE females, and then crossing the resulting *Polg*^wt/D257A^TwOE to one another. The Mutator *Polg*^D257A^ allele was maintained through the paternal line to avoid the inheritance of mtDNA mutations, and only present in females of the last cross when necessary to obtain homozygote *Polg* ^D257A/D257A^ offspring.

Adult mice were euthanized by gradual inhalation of excess CO_2_ and younger mice (< P7) by cervical dislocation. Tissues were weighed and collected immediately, snap-frozen in liquid nitrogen and stored at -80°C or histologically processed as detailed below.

Mouse embryonic fibroblasts (MEFs) were extracted from E16.5 embryos using standard procedures: dissociation of the embryo body (without head, heart and liver) using a scalpel and followed by incubation in trypsin:EDTA, plating in gelatine-coated plates and growth in standard MEF media.

The animal work was performed according to regulations and approval of The Ethical Review Board of Finland (permit: ESAVI/3686/2021).

### Experimental design

The experimental cohort was generated as detailed above. Each mouse was considered an independent biological replicate (n), and the experimentation largely consisted in comparing different genotypes and their littermate controls. Due to the limited amount of biological material (e.g. minute neonatal hearts), many methods required the entirety of the sample and thus the n number varies across measurements, ensuring the statistical power and optimizing the use of the material.

### DNA isolation

DNA was extracted from frozen samples by standard phenol-chloroform extraction. For extraction of DNA from paraffinized samples, 6-7 slices of paraffin were cut from the block. Paraffin was removed by washing the slides twice with xylene and the samples were rehydrated in a descending ethanol series (100%, 90%, 75%). The samples were dried and the extraction was continued according to the standard protocol, adding a 20-minute incubation at 90°C to de-crosslink DNA at the end of the proteinase K incubation.

### RNA isolation and RT-qPCR

RNA was extracted using Trizol reagent (Invitrogen) and purified through Qiagen RNeasy minicolumns (Qiagen), following the manufacturer’s instructions.

For RT-qPCR, cDNA was synthesized with Maxima First Strand cDNA Synthesis Kit (Thermo Scientific) according to the manufacturer’s instructions. 100 ng of RNA was used in the cDNA synthesis and the cDNA was diluted 1:4 before its use in RT-qPCR. RT-qPCR was performed with SensiFAST^™^ SYBR No-ROX Kit (BIO-98020, Bioline) and the relative gene expression levels were normalized against β-actin expression. The primers used are listed in the supplementary material.

### mtDNA quantitation

MtDNA copy number was quantified by quantitative PCR (qPCR) using the SensiFAST^™^ SYBR No-ROX Kit (BIO-98020, Bioline), by amplification of three mtDNA fragments (12S-rRNA, COX1 and ND4) against nuclear-DNA-encoded RBM15. 25 ng of DNA was used per PCR reaction, all qPCR assays were run in triplicates. The primers used are listed in the supplementary material.

### Histology & immunofluorescence

Formalin-fixed paraffin-embedded tissues or frozen sections were used for histological stainings and histochemical activity analyses. For immunohistology, standard protocols were used with slight modifications together with manufacturer instructions. Antigen retrieval from paraffin-derived samples was done in 10Mm citric acid buffer pH 6,0. The antibodies used are found in the supplementary material. Frozen sections were used for the laminin immunostaining and the OilRedO staining.

For immunofluorescence, coverslips with cultured cells were fixed in 4% PFA at room temperature for 15 min and washed with PBS. Slides were blocked in blocking buffer (1% BSA, 0.1% TritonX and 10% horse serum in PBS), followed by overnight primary antibody incubation in antibody buffer (1% BSA, 0.1% TritonX and 1% horse serum in PBS). Following PBS washing, samples were incubated for 1 hour at room temperature in secondary antibodies conjugated with Alexa Fluor fluorescent probes (Thermo Fisher Scientific) diluted 1:400 in 1% PBS. Finally, the coverslips were mounted onto slides with DAPI-containing medium (Vectashield #H-1200-10).

### EdU mtDNA replication assay

EdU incorporation into nucleoids was performed using the Click-iT^™^ EdU Alexa Fluor^™^ 488 Imaging Kit (Fisher Scientific), following the manufacturer’s instructions with minor deviations (non-confluent cells incubated for 1h in 50 μM EdU-containing media, and two sequential Click reactions to ensure detection). The resulting fixed cells were blocked and immunostained with a dsDNA antibody (details in supplementary material) and DAPI, to identify nucleus and mtDNA nuclei as above.

### Microscopy and image analysis

For transmission electron microscopy, fresh heart samples were fixed in 2.5% glutaraldehyde (EM-grade) in 0.1 M sodium phosphate buffer for 2 h, RT, and transferred into 0.1 M sodium phosphate buffer (pH 7.4) after fixation. The samples were postfixed with 1% reduced osmium tetroxide in 0.1 M sodium phosphate buffer for 1 h, on ice. Samples were again washed with phosphate buffer and then dehydrated through a series of ethanol and acetone prior to gradual infiltration into Epon (TAAB 812, Aldermaston, UK). The resin was polymerized at 60°C for a minimum of 16 hours. A pyramid was trimmed on the tissue and 60-70 nm sections were cut with an ultramicrotome and placed on Pioloform-coated copper grids. Sections were post-stained with uranyl acetate and lead citrate. TEM micrographs were acquired using a JEM-1400 transmission electron microscope (Jeol Ltd., Tokyo, Japan) running at 80 kV, with a bottom-mounted CCD camera (Orius SC 1000B, AMETEK-Gatan Inc., Pleasanton, CA), with images of 4008 × 2672 pixels.

Fluorescent images were acquired using the Andor Dragonfly spinning disk confocal microscope, and image quantification was performed with the CellProfiler software, using its standard segmentation and classification modules.

### Echocardiography

To analyze cardiac function and dimensions of the left ventricular chamber, P7 pups were anesthetized by inhalation with 2,5% isoflurane mixed with 0.5L/min 100% oxygen (Vevo Compact Dual Anesthesia System). The mice were placed on a heating stage (Vevo Imaging Station) in supine position and 2,5% isoflurane mixed with 0.5L/min 100% oxygen was supplied continuously via a nose cone to maintain the mice under sedation. Pre-warmed ultrasound gel was applied to the thoracic area and two-dimensional ECG images were acquired using MS550D 22-50 MHz linear array solid-state transducer (Vevo 2100 Ultrasound, FUJIFILM VisualSonics). Long-axis ECG images in B-mode were used to confirm the anatomic boundaries of the ventricular chambers. M-mode images along the parasternal short axis were used to measure left ventricular internal diameter, left ventricular posterior wall thickness, and interventricular septum thickness at end-systole and end-diastole. These parameters were used to calculate left ventricular mass, volume, ejection fraction, and fractional shortening (Vevo Vasc Analysis software).

### MtDNA sequencing

mtDNA was enriched from whole P7 heart DNA by long-range PCR amplification of three fragments; processed and analyzed as in ^55^. DNA concentrations were measured with Qubit and quality was assessed using Genomic DNA ScreenTape Assay (Agilent TapeStation 4200). 100 ng of pooled amplicon DNA (DIN > 6.8) was converted into sequencing libraries using NEBNext Ultra II FS DNA Library Prep Kit for Illumina. Completed libraries were pooled and then sequenced with a NextSeq Mid Output 300 cycle flow cell on the NextSeq 500 to produce 2×150 bp reads.

### MtDNA topology

The analysis of mtDNA topology was performed as in ^56^, 1 μg of total DNA was separated over a 0,4 % agarose gel (UltraPure Agarose, # 16500, Life Technologies) without Ethidium bromide in 1x TBE buffer overnight at 32 V. The gel was treated for 2x 15 min with 0,25 M HCl and 2x 20 min with 0,5 M NaOH, 1,5 M NaCl, transferred by capillary blot onto Hybond XL-nylon membrane and hybridized in Church’s buffer with a cytochrome B probe (nucleotides 14783-15333 of mouse mtDNA).

### RNA sequencing

The sequencing services were provided by the Biomedicum Functional Genomics Unit at the Helsinki Institute of Life Science and Biocenter Finland at the University of Helsinki.

3’UTR RNA sequencing “Bulkseq” was performed as in ^57^ using whole E16.5 heart RNA (isolated as described above). Shortly, Bulkseq involves mRNA priming with an oligo dT primer which also contains a 12 bp sample barcode and an 8 bp UMI sequence which can be used to remove PCR duplicates during data analysis. Single-stranded cDNA is then converted to double-stranded cDNA using the template switch effect and the double-stranded cDNA product is then amplified with PCR and another set of primers (SMART PCR primer). Samples are then pooled together, and PCR sequencing pools are made with Nextera i7 primers and the Dropseq P5 primer. Sequencing is then performed on the NextSeq High Output 75 cycle flow cell on the NextSeq 500 to produce (Read 1: 20 bp, Index 1 (i7): 6 bp and Read 2 (61 bp)).

For data analysis, bcl2fastq2 was used to convert BCL files to FASTQ file format and demultiplex the samples. Sequenced reads were trimmed for adaptor sequence and reads shorter than 20nt were also removed using Trimmomatic. Read processing was performed using drop-seq tools (version 2.4.0); In short, reads were additionally filtered to remove polyA tails of length 6 or greater and then trimmed reads were aligned to GRCm38 reference genome (GENCODE Mouse Release 25) using STAR aligner (2.7.6a). Gene expression was quantified with drop-seq tools after removing PCR duplicates using unique molecular identifier -sequences (UMIs).

Differential expression analysis was done in the DESeq2 software in the R environment. The count values were normalized between samples using a geometric mean. Sample-wise factors were estimated to correct for library size variability and estimation of dispersion (i.e. variance, scatter) of gene-wise values between the conditions. A negative binomial linear model and Wald test were used to produce p-values. Low-expression outliers were removed using Cook’s distance to optimize the p-value adjustment and finally, multiple testing adjustment of p-values was done with the Benjamini-Hochberg procedure.

### Targeted metabolomics profiling analysis

Metabolites were extracted from 6-10mg P1 mouse heart tissue using a 2 mL Precellys homogenization tube (Bertin Technologies, Montigny-le-Bretonneux, France) with 2.8 mm ceramic (zirconium oxide) beads with 400 µL of cold extraction solvent (Acetonitrile:Methanol:Milli-Q Water; 40:40:20). Subsequently, samples were homogenized using a tissue homogenizer (Bertin Technologies, France) for 3 cycles (30sec at 5500rpm with 60sec pause at 4°C). Followed by centrifugation at 14000rpm at 4°C for 5 minutes. The supernatant was loaded into a Phenomenex, Phree Phospholipid removal 96 well plate 30mg (Part No. 8E-S133-TGB) and passed through using a robotic vacuum. The filtrate was transferred into polypropylene tubes and placed into a Nitrogen gas evaporator to dry the solvent completely. Dried samples were suspended with 40µL of mobile phase solvent (Acetonitrile: 20mm Ammonium Hydrogen Carbonate Buffer, pH 9.4; 80:20) and vortex for 2 minutes and transfer into HPLC glass autosampler vials. 2µL of sample injected with Thermo Vanquish UHPLC coupled with Q-Exactive Orbitrap quadrupole mass spectrometer equipped with a heated electrospray ionization (H-ESI) source probe (Thermo Fischer Scientific). A SeQuant ZIC-pHILIC (2.1×100 mm, 5-μm particle) column (Merck) used for chromatographic separation. Gradient elution was carried out with a flow rate of 0.100 mL/minutes using 20mM ammonium hydrogen carbonate, adjusted to pH 9.4 with ammonium solution (25%) as mobile phase A and acetonitrile as mobile phase B. The gradient elution was initiated from 20% mobile phase A and 80% of mobile phase B and maintained till 2min. After that, 20% mobile phase A gradually increase up to 80% till 17 min, then 80% to 20% Mobile phase A decrease in 17.1 min and maintained up to 24 minutes. The column oven and auto-sampler temperatures were set to 40 ± 3 °C and 5 ± 3 °C, respectively. MS equipped with a heated electrospray ionization (H-ESI) source using polarity switching and the following setting: resolution of 35,000, the spray voltages: 4250 V for positive and 3250 V for negative mode, the sheath gas: 25 arbitrary units (AU), and the auxiliary gas: 15 AU, sweep gas flow 0, Capillary temperature: 275°C, S-lens RF level: 50.0. Instrument control operated with the Xcalibur 4.1.31.9 software (M/s Thermo Fischer Scientific, Waltham, MA, USA).

In the data processing, the final peak integration was done with the TraceFinder 4.1 software (Thermo Fischer Scientific) using confirmed retention times of 462 metabolites in-house library developed using library kit MSMLS-1EA (Merck). For further data analysis, the peak area data were exported as an excel file. Data quality was monitored throughout the run using a pooled sample as Quality Control (QC) prepared by pooling 5µL from each suspended sample and interspersed throughout the run as every 10th sample. After integration of QC data with TraceFinfer 4.1 detected metabolites were checked for peak, % RSD were calculated, and acceptance limit was set ≤ 20%. Blank samples for carryover were injected after every fifth randomized sample to monitor the metabolites’ carryover effect and calculated against the mean QC area and the acceptance limit was set ≤ 20% for each metabolite. Background % noise was calculated with respect to the first blank against the mean QC area and the acceptance limit was set ≤ 20% for each metabolite.

The obtained data were analyzed and explored using the MetaboAnalyst software following the recommended path. The data was normalized by the sum of the sample, Generalized log transformation (glog2) and Autoscaling (mean-centered and divided by the standard deviation of each variable) settings were used.

### Proteomic analysis

The mass spectrometry analyses proteomic were performed at the Turku Proteomics Facility supported by Biocenter Finland, following their pipeline. Shortly, proteins were isolated from approximately five (5) mg of snap-frozen hearts pulverized with polypropylene grinding rods in liquid nitrogen, extracted with lysis buffer (6 M Urea, 100 mM TEAB and 1 x Halt protease inhibitor cocktail) and sonication (3 × 5 min cycles of 30s on/off; Diagenode Bioruptor sonication device). Extracted proteins protein were digested to peptides with in-solution digestion performed at the Turku Proteomics Facility, desalted with a Sep-Pak C18 96-well plate (Waters) and evaporated to dryness and stored at -20 0C.

Digested peptide samples were dissolved in 0.1% formic acid and peptide concentration was determined with a NanoDrop device. For DIA analysis 800 ng peptides was injected and analyzed in random order. Wash runs were submitted between each sample to reduce the potential carry-over of peptides. The LC-ESI-MS/MS analysis was performed on a nanoflow HPLC system (Easy-nLC1200, Thermo Fisher Scientific) coupled to the Orbitrap Lumos Fusion mass spectrometer (Thermo Fisher Scientific) equipped with FAIMS interface and a nano-electrospray ionization source. Peptides were first loaded on a trapping column and subsequently separated inline on a 15 cm C18 column (75 μm x 15 cm, ReproSil-Pur 3 μm 120 Å C18-AQ, Dr. Maisch HPLC GmbH, Ammerbuch-Entringen, Germany). The mobile phase consisted of water with 0.1% formic acid (solvent A) or acetonitrile/water (80:20 (v/v)) with 0.1% formic acid (solvent B). A 110 min gradient was used to elute peptides (70 min from 5% to 21% solvent B followed by 40 min from 21 % to 36 min solvent B). Samples were analyzed by a data-independent acquisition (DIA) LC-MS/MS method. MS data were acquired automatically by using Thermo Xcalibur 4.1 software (Thermo Fisher Scientific). In a FAIMS-DIA method, a duty cycle contained two compensation voltages (−50V and -70V) with one full scan (400-1000 m/z) and 30 DIA MS/MS scans covering the mass range 400 -1000 with variable width isolation windows in each of the compensation voltages. Data analysis consisted of protein identifications and label-free quantifications of protein abundances. Data were analyzed by Spectronaut software (Biognosys; version 15.0.2). DirectDIA approach was used to identify proteins and label-free quantifications were performed with MaxLFQ. Main data analysis parameters in Spectronaut included - Enzyme: Trypsin/P, Missed cleavages: 2, Fixed modifications: Carbamidomethyl, Variable modifications: Acetyl (protein N-term) and oxidation (M), Protein database: Swiss-Prot 2021_02 Mus Musculus, Normalization: Global median normalization, Differential abundance testing: Unpaired t-test and multiple-testing correction of the p-values with Benjamini-Hochberg method (Q-values).

## Supplemental information, figures and legends

Supplementary data including the detailed results of the omics analyses, numerical data used for graphical representation in the figures and representative videos of echocardiograms can be found at https://doi.org/10.17632/wm2dhwpghm.1 ^58^.

**Supplementary figure 1**. (A) Fat accumulation in the P7 liver; OilRedO staining. (B) Disrupted alveolar ultrastructure; hematoxylin/eosin staining of the lung. (C) Relative mtDNA copy number to nuclear gene RBM15, using three different mtDNA sites (genes for 16S rRNA, COXI & ND4) in P7 liver and lung. (D-E) Blood hemoglobin and glucose concentrations at P7. P values from a two-tailed t-test as stars: * (p < 0.05), ** (p < 0.01), *** (p < 0.001). (F) Nuclear DNA integrity in stem cells: In testes and skin epidermis, gH2AX (marking DNA double-strand breaks) in cyan, PCNA (actively replicating nuclei) in magenta, and keratin-14 (marking the basal layer of the skin) in yellow. Positive controls from 8-month-old mutator (testes) and DNA-repair deficient embryo (skin). (G) Immunofluorescent dsDNA staining of P7 hearts, showing enlarged nucleoids (white arrowhead). (H and I) Characterization of linear mtDNA fragments accumulating in Mut heart mtDNA. (H) P7 heart DNA digested with DraI and probed with nucleotides 14783-15333 of mtDNA. The sketch visualizes the linear fragments arising from intact mtDNA as well as mtDNA having strand breaks close to OH. (I) P7 heart DNA digested with SacI and probed with nucleotides 5399-5997 of mtDNA.

**Supplementary figure 2**. (A) Volcano plots depicting the fold-change and statistical significance of proteomic changes between WT and Polg^Mut^TwOE groups, classified by their top KEGG pathway. The dotted line indicates the significance threshold at Q < 0.01. Arrowheads indicate the position of cropped data points. (B) Log2-fold change between WT and Polg^Mut^TwOE groups of the detected proteinogenic amino acids. Q values as stars: * (p < 0.05), ** (p < 0.01), *** (p < 0.001).

