## Supplementary figures and images for "Overactive mitochondrial DNA replisome causes neonatal heart failure via ferroptosis"

### Supplementary figure 1

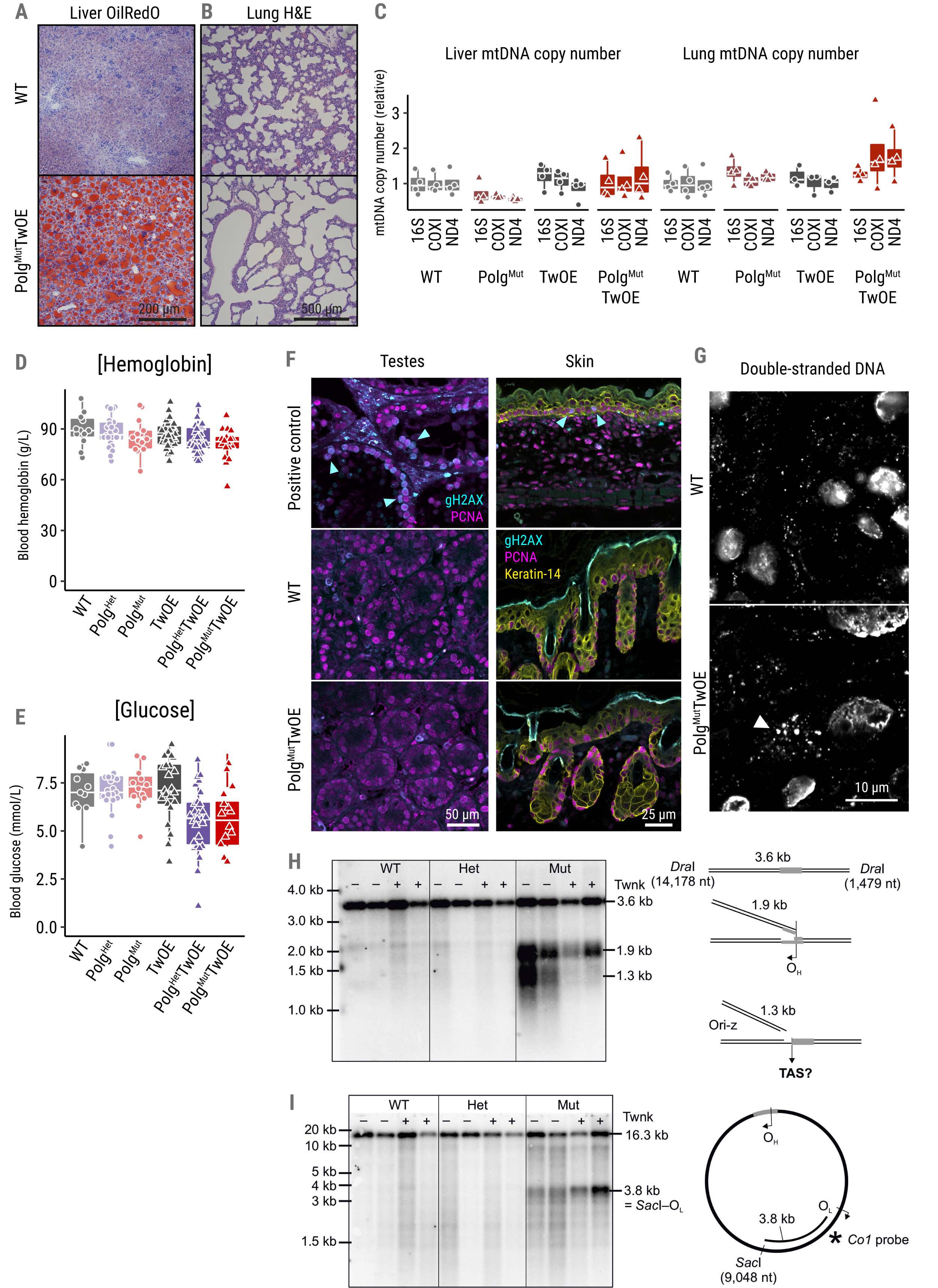

### Supplementary figure 2

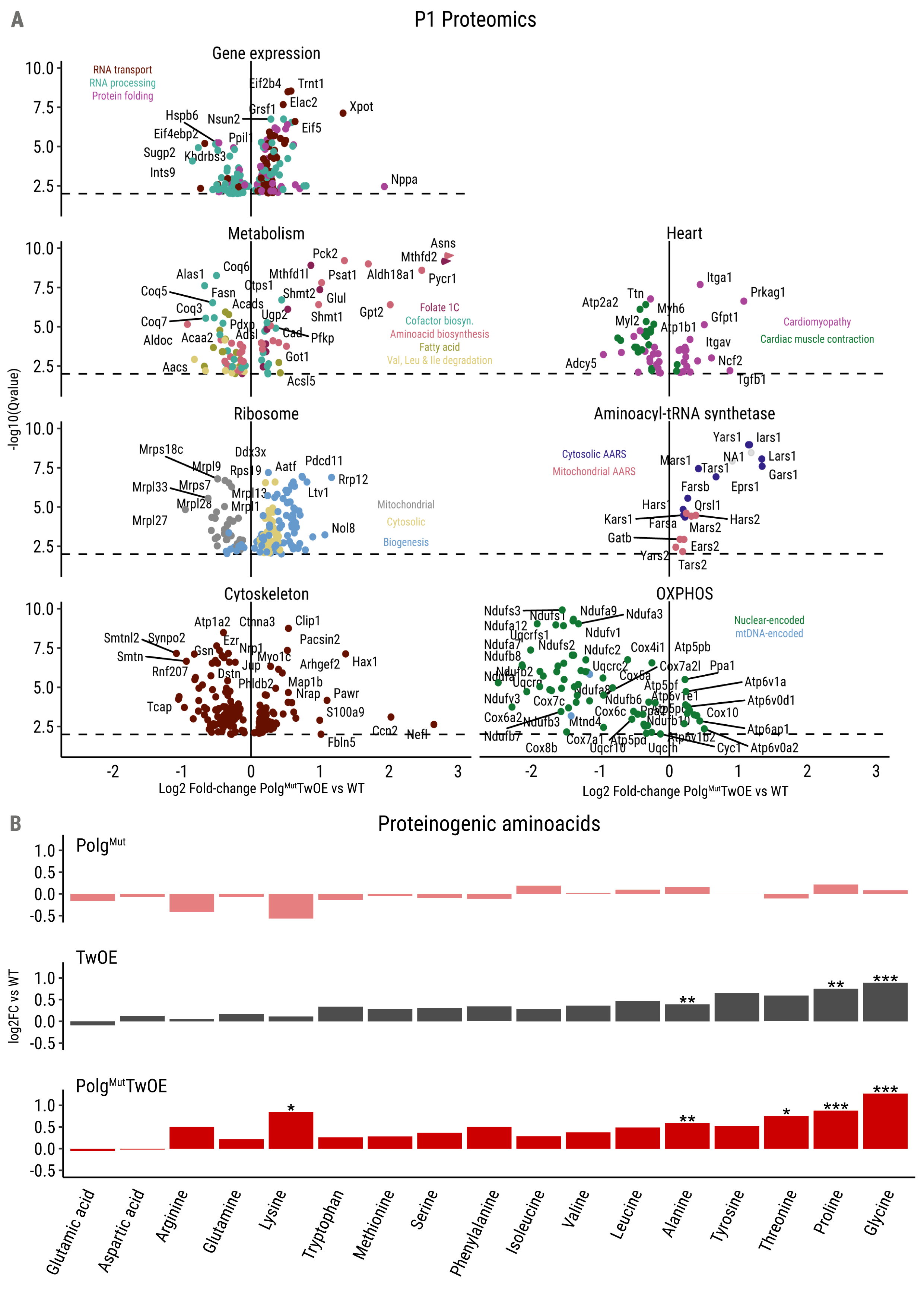
